# Geometrical confinement guides Brachyury self-patterning in embryonic stem cells

**DOI:** 10.1101/138354

**Authors:** Blin Guillaume, Catherine Picart, Manuel Thery, Michel Puceat

**Affiliations:** MRC Centre for Regenerative Medicine, Institute for Stem Cell Research, School of Biological Sciences, University of Edinburgh, Edinburgh, United Kingdom; Université de Grenoble Alpes, Grenoble Institute of Technology, CNRS, UMR 5628, LMGP, 3 parvis Louis Néel, F-38016, Grenoble, France; CytoMorpho Lab, Biosciences & Biotechnology Institute of Grenoble, UMR5168, CEA/INRA/CNRS/Université Grenoble-Alpes, Grenoble, France; CytoMorpho Lab, Hopital Saint Louis, Institut Universitaire d’Hematologie, UMRS1160, INSERM/Université Paris Diderot, Paris, France; Faculté de Médecine La Timone, Université Aix-Marseille, INSERM UMR_910, Marseille, France

## Abstract

During embryogenesis, signaling molecules initiate cell diversification, sometimes via stochastic processes, other times via the formation of long range gradients of activity which pattern entire fields of cells. Such mechanisms are not insensitive to noise (Lander, 2011), yet embryogenesis is a remarkably robust process suggesting that multiple layers of regulations secure patterning during development. In the present study, we present a proof of concept according to which an asymmetric pattern of gene expression obtained from a spatially disorganised population of cells can be guided by the geometry of the environment in a reproducible and robust manner. We used ESC as a model system whithin which multiple developmental cell states coexist (MacArthur and Lemischka, 2013; Smith, 2017; Torres-Padilla and Chambers, 2014). We first present evidence that a reciprocal regulation of genes involved in the establishment of antero-posterior polarity during peri-implantation stages of mouse development is spontaneously occuring within ESC. We then show that a population of cells with primitive streak characteristics localise in regions of high curvature and low cell density. Finally, we show that this patterning did not depend on self-organised gradients of morphogen activity but instead could be attributed to positional rearrangements. Our findings unveil a novel role for tissue geometry in guiding the self-patterning of primitive streak cells and provide a framework to further refine our understanding of symmetry breaking events occuring in ESC aggregates. Finally, this work demonstrates that the self-patterning of a specific population of ESC, Brachyury positive cells in this case, can be directed by providing engineered external geometrical cues.

## Introduction

Developmental patterning is the process through which spatially defined regions of distinct cell types emerge from a group of cells initially seemingly equivalent. During early embryonic development, such a process requires a symmetry breaking event in order to generate the first landmarks which will define the future axes of the body. In many species, it is well recognised that maternal determinants or fertilisation define this break of symmetry (Gilbert, 2013). In mammals however, due to the regulative nature of the embryo, the origin of the first asymmetries remains intensively debated (Arnold and Robertson, 2009; Rossant and Tam, 2009; Zernicka-Goetz et al., 2009). In the mouse, antero-posterior (AP) polarity becomes apparent during the peri-implantation stages (Fig. 1A). During this period, the embryo adopts an elongated shape described as an egg-cylinder. The pluripotent cells of the epiblast are flanked by the extraembryonic ectoderm (ExE) on the proximal side of the embryo and are sitting on a layer of extraembryonic epithelial cells called the visceral endoderm (VE). The AP axis emerges when subsets of the VE cells adopt distinct molecular signatures. The proximo-posterior side of the embryo is marked by the expression of Wnt3 (Rivera-Pérez and Magnuson, 2005) which engage in a signaling autoregulatory loop involving Nodal from the epiblast and BMP4 from the ExE (Ben-Haim et al., 2006; Brennan et al., 2001). Nodal and BMP4 participate in the specialisation of distal VE (DVE) cells (Kimura-Yoshida et al., 2005; Rodriguez et al., 2005; Yamamoto et al., 2004) which subsequently migrate towards the anterior side to become the anterior VE (AVE) (Ding et al., 1998; Rodriguez et al., 2005; Srinivas et al., 2004) (reviewed in (Stower and Srinivas, 2014)). The AVE, composed of multiple cell subpopulations, secretes antagonists of Nodal, BMP and Wnt such as Cerberus, Lefty1 or Dkk1, thus constituting a negative feedback which restricts the activity of Nodal/Wnt/BMP to the posterior side of the embryo (Belo et al., 1997; Glinka et al., 1998; Kimura-Yoshida et al., 2005; Meno et al., 1996; Yamamoto et al., 2004). Gastrulation becomes apparent at around E6.5 with the formation, under Wnt3 influence (Barrow et al., 2007; Liu et al., 1999; Yoon et al., 2015), of the primitive streak (PS) on the proximo-posterior side of the epiblast. The PS is characterised by the expression of early mesendodermal markers such as T-Brachyury (T) (Beddington et al., 1992; Wilkinson et al., 1990), loss of epithelial characteristics reviewed in (Morali et al., 2013) and an inversion of polarity prior migration of the ingressing cells (Burute et al., 2017; STERN, 1982). Importantly, it is the former positioning of extraembryonic structures, i.e. Wnt3+ VE cells and the AVE, which define AP polarity and which enable the establishment of an AP gradient of signaling molecules which regionalises the epiblast.

**Fig. 1.**
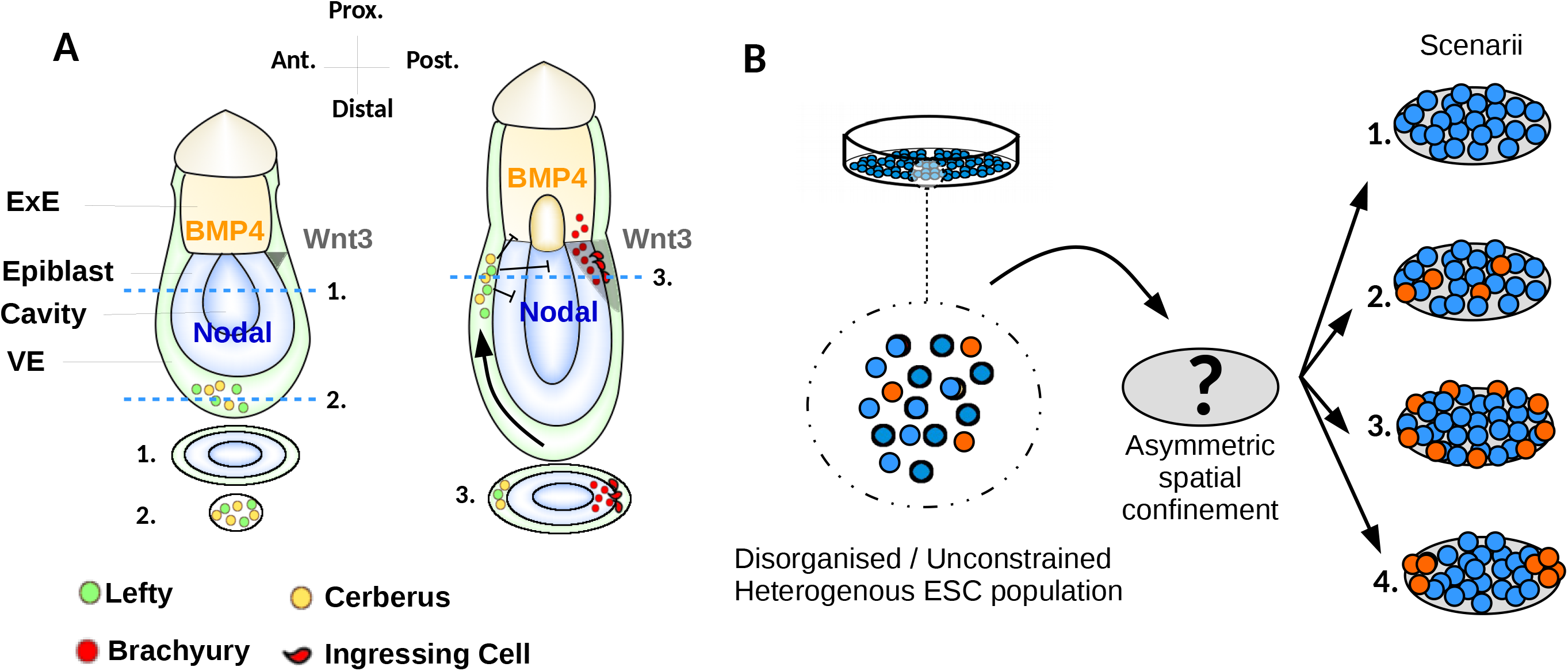
Embryonic symmetry breaking event in vivo, a possible role for geometrical constraints in guiding the axis. **A.** This schematic illustrates the establishment of AP polarity during the peri-implantation stages of mouse development (described in the introduction). **B.** Methodological approach and tested hypotheses. ESC in culture contain subpopulations with disting genetic profiles. Spatial confinement may 1) regulate gene expression, 2) have no apparent effect, 3) enable patterning via border effects in a symmetry insensitive fashion, 4) enable patterning with geometry guiding spatial organisation.

Embryonic stem cells (ESC) are cell lines derived from the pluripotent cells of the pre-implantation embryo (Evans and Kaufman, 1981; Martin, 1981). When reintroduced into a blastocyst, ESC can contribute to the epiblast to paticipate in the making of a new organism (Bradley et al., 1984). On the other hand, ESC are very inefficient in integrating into extraembryonic lineages (Beddington and Robertson, 1989). Yet we emphasized earlier that regionalisation of extraembryonic lineages precedes and controls the regionalisation of the epiblast. For these reasons, AP patterning has long been thought to be a prerogative of embryonic development. However the emergence of embryo-looking AP patterning events have been recently reported to occur within 3D aggregates of pluripotent cells in culture (ten Berge et al., 2008; Brink et al., 2014; Harrison et al., 2017; Marikawa et al., 2009), revealing the possibility to reanimate in vitro the stunning self-organising competence of these cells observed in vivo.

These remarkable findings call to mind how the questions surrounding early embryonic patterning may be formulated in engineering terms (Davies, 2017; Sasai, 2013). Indeed, an interesting approach is to ask which is the minimal set of external instructions to be imposed to allow ESC to recapitulate a normal developmental patterning program.

In the absence of exogenous signaling molecules, ESC homogeneously adopt an anterior neural fate (Eiraku et al., 2011; Meinhardt et al., 2014; Ying et al., 2003a). This fate can be antagonised with BMP signaling which in the embryo is delivered by the ExE. The strategy of Harrison and colleagues has been to assemble ESC with trophoblast stem cells (an in vitro counterparts of the ExE) in order to provide a localised source of BMP4 (Harrison et al., 2017). This coculture system embedded in matrigel, self-organised into a structure which closely resembled the egg cylinder stage embryo including the apparition of T+ cells preferentially localised to one side of the edifice.

In other studies, the polarised pattern of gene expression in differentiating pluripotent cell aggregates have been reported even in the absence of a localised source of signaling (ten Berge et al., 2008; Marikawa et al., 2009), demonstrating that symmetry breaking is a competence which is inherent to ESC. Symmetry breaking events in such aggregates may become a highly reproducible process if specific requirements are met. These include a serum-free environment, canonical wnt activation, 3D aggregation and a precise starting number of cells (Brink et al., 2014).

The studies mentionned above are in line with the work from several other groups who reported the spectacular degree of auto-organisation emanating from 3-dimensional culture of differentiating pluripotent stem cells (so-called organoids) with no requirement for a supply of a localised signal (Eiraku et al., 2011; Lancaster et al., 2013; Meinhardt et al., 2014; Nakano et al., 2012). If 3-dimensionality reveals ESC self-organising abilities, 2D micropatterning techniques have already proven useful to understand the forces at work. Pioneering studies with ESC (Bauwens et al., 2008; Davey and Zandstra, 2006; Peerani et al., 2007, 2009) and with multipotent cells (McBeath et al., 2004) have shown that spatial confinement of colonies of cells on 2D patterns allow to harness and challenge the environment sensing abilities of cells in culture. What these works have especially demonstrated is the ability of stem cells to form their own niche, i.e. to generate their own gradients of morphogens and their competence to interpret signals in a position dependent manner.

Together with the identification of the molecular factors involved during early development, these founding works have paved the way to the recent establishment of a method to recapitulate several aspects of the early gastrulating embryo with human ESC (Etoc et al., 2016; Warmflash et al., 2014). Using disc shaped micropatterns to provide spatial confinements and BMP4 to trigger differentiation, the method enables the formation of spatially defined domains of cells with ectodermal, mesodermal, endodermal and trophectodermal characteristics. Recently, the size of the patterns and the concentration of the initial inducing signal was systematically tested. This has led to further insights into the nature of interactions between diffusible signals and emergent lineages in the culture to provide a model of self-organisation which can scale with size (Tewary et al., 2017).

Common principles have emerged from the studies mentioned above. In order to allow spontaneous patterning to become apparent in ESC culture: 1) The presence in the medium of a signaling molecule which antagonises anterior-neural fate such as BMP4 or Activin is required to prevent all cells from turning into neurectodermal cells. 2)This molecule does not need to be provided in a localised manner if a 3D culture is employed or if spatial confinement is provided to enable endogenous self-organising mechanisms to take place. 3) The length scale of the confinement or the size of the starting population has drastic effects on the outcome of the process and needs to be optimised in each case (Bauwens et al., 2008; Brink et al., 2014; Tewary et al., 2017)

The delineation of the constraints on cell signaling and cell number required to observe patterning within in vitro cultures provides a great deal of information about the mechanisms at work. One important question which has not yet been addressed is whether the axis of an autonomous self-patterning event is sensitive to geometrical constraints and thus can be guided with engineered extrinsic cues. In the present work we interrogate the possibility of asymmetric geometrical confinement to fulfil such a role (Fig. 1B). Using micropatterns, we demonstrate that geometry predicts the preferential spatial organisation of PS progenitors in vitro. Using pharmacological compounds and RNA interference to manipulate both autonomous morphogen gradients and cell positioning within differentiating ESC colonies, we further describe how DVE and PS markers expressed in the culture depend on autonomous paracrine signaling. We provide evidence that endogenous Nodal and Wnt3 define the number of primitive streak progenitors in the population but not their localisation, which appears to depend instead on group geometry which guides cell movements within the colony. We discuss the implications of these findings for pattern formation in ESC aggregates and during gastrulation.

## Results

### Proximal and distal gene expression within ESC culture

ESC culture can be seen as a dynamic equilibrium of multiple developmental states (MacArthur and Lemischka, 2013; Torres-Padilla and Chambers, 2014). Since ESC generate their own niche in culture and since they are responsive to local changes in cell density (Davey and Zandstra, 2006), we decided to first investigate if early developmental markers would respond to changes in cell density and if so, how they would be regulated.

We plated cells on gelatin-coated dishes in FCS and LIF containing medium. We seeded cells at low density (5000 cells/cm2), high density (50 000 cells/cm2) or at an optimal density for ESC propagation (12 000 cells/cm2) referred thereafter as control density (Fig. 2A). Additionally we developed a method of micropatterning to force the cells to grow as colonies of defined shape and size in order to uncouple global from local cell density. In this experiment, we chose to culture cells on discoidal µP with a diameter of 195µm (30000µm^2^), and with µP centers separated by 490µm (Fig. 2A). Colony sizes >100µm was shown to maximize signaling pathways such as Stat3 activation and separation of >400µm was found to be the minimal distance to consider colonies independent for this same pathway (Peerani et al., 2009). This configuration permitted the total cell number per dish to be equivalent to the total number of cells in the control density while promoting a local cell density which would normally be found at high density. We cultured the cells for 2 days in LIF containing medium. We first monitored by qPCR the level of expression of pluripotency markers Oct4, Sox2, Nanog and Rex1. We observed only minor changes between the different conditions (Fig. 2B). Only *Nanog* was increased twofold and *Oct-4* decreased twofold at high cell density compared to the control condition.

**Fig. 2.**
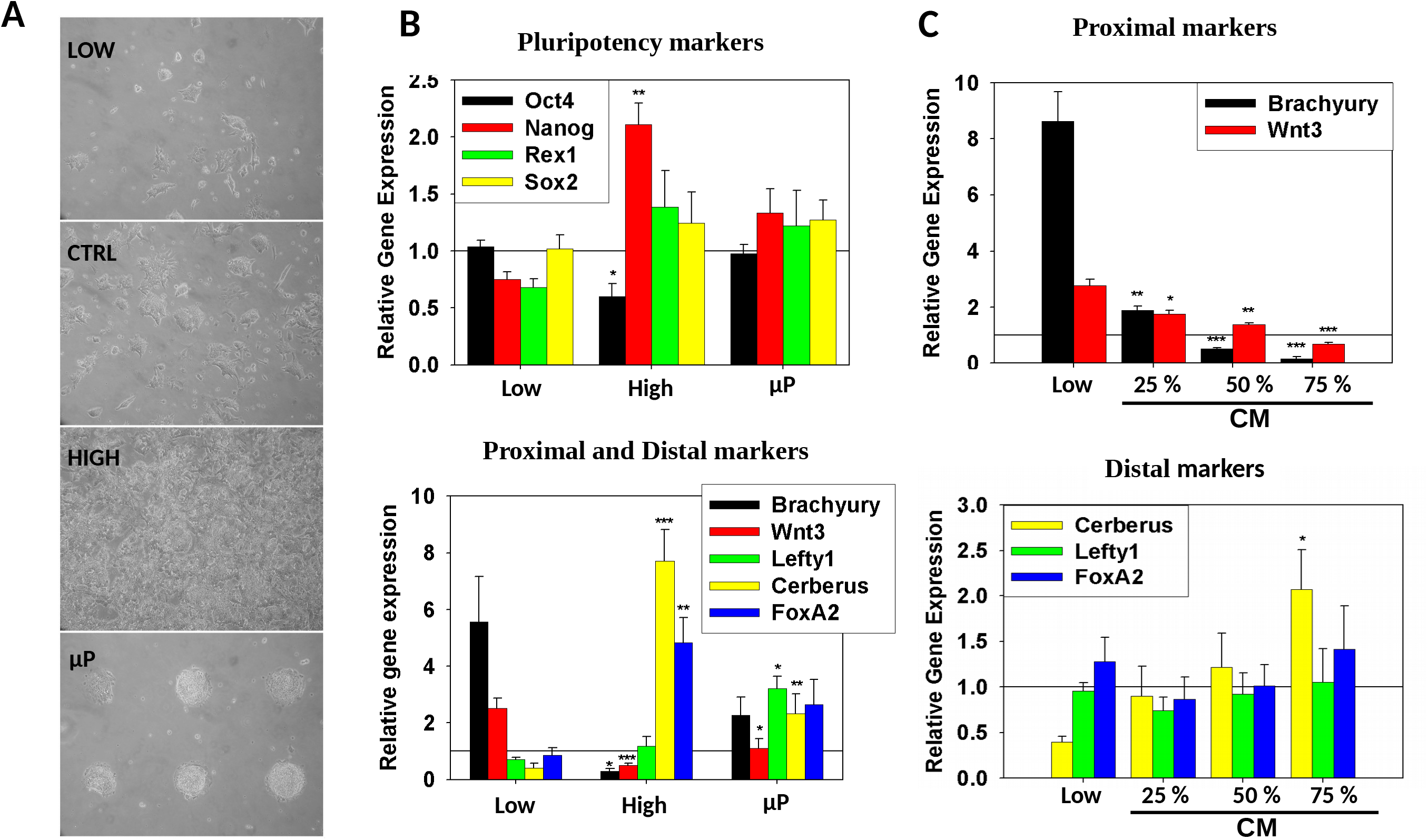
A balance of proximal and distal embryonic markers exist in ESC culture and is regulated by diffusible molecules. **A.** Phase contrast micrographs of ESC cultures at control (12 000 cells/cm2), low (5000 cells/cm2) or high density (50 000 cells/cm2) or on µP, scale bar 100µm. **B.** Gene expression profile of cells grown at different densities or on µP. Pluripotency genes (upper panel) and early asymmetric markers (lower panel). **C.** Gene expression profile of cells grown at low density with increasing amount of conditionned medium (CM). Real-time PCR data have been normalised to the control density (reference line) in each experiment. Error bars represent the S.E.M of at least 3 independent experiments. The stars represent the p value of the Student’s t-test as compared to low density (* p < 0.05, ** p < 0.01, ***p < 0.005).

Next, we decided to monitor the expression of early developmental genes. The promoter activity of several pro-differentiation factors have been reported in ESC culture and have been shown to mark subpopulations of cells with biases towards specific routes of differentiation (Canham et al., 2010; Davies et al., 2013; Niakan et al., 2013; Singh et al., 2007). Interestingly, most of these genes are expressed around peri-implantation stages before gastrulation. We thus decided to investigate the regulation of genes which are expressed before or at the onset of gastrulation. We found that the levels of the proximal markers, Brachyury (T) and Wnt3 (Rivera-Pérez and Magnuson, 2005), were negatively correlated to cell density. Conversely, the AVE markers Cerberus (Belo et al., 1997) and FoxA2 (Kimura-Yoshida et al., 2007) were upregulated at high density and downregulated at low density. These results suggest that low density promotes proximo-posterior identity in ESC whereas high density favours an environment permissive for anterior lineages.

Interestingly, we observed a reproducible trend of increase of T, Cerberus and FoxA2 on micropatterns (µP) as compared to the control density (Fig. 2B bottom panel). This indicates that on µP, the two cellular contexts which favour the expression of these genes may coexist. Surprisingly we also observed Lefty1 to be upregulated on uP while density did not affect this gene. This observation may indicate that cells grown on patterns become more responsive to Nodal/Activin activity since Lefty1 is a known direct target of the smad2/3 pathway (Besser, 2004).

To investigate which effect could be attributed to paracrine signals and which one to cell/cell contacts or local constraints, we used conditioned medium (CM) from cells grown at high density for two days and tested the effect of several ratios of this CM and fresh medium onto cells cultured at low density for two days (Fig. 2C). Under these experimental conditions, T and Wnt3 downregulation as well as Cerberus upregulation correlated with the increasing proportion of CM. This result demonstrates that diffusible signaling molecules are involved in the regulation of these genes. Conversely, Lefty1 and FoxA2 expressions were not affected by the addition of CM indicating that local mechanical cues or juxtacrine may be required to modulate their expression.

Together, these results support the idea that ESCs in culture recapitulate the reciprocal regulation of proximal and anterior genes observed in vivo during the establishment of AP polarity.

T has been widely used to identify PS formation in mice or in ESC (Gadue et al., 2006; Turner et al., 2014a). However, it has also been shown to be expressed initially in the ExE in vivo by ISH(Rivera-Pérez and Magnuson, 2005). For this reason, it was suggested that caution should be taken when using T as a PS marker since the PS is derived from epiblast cells. As ESC are known to poorly differentiate into the ExE lineage (Beddington and Robertson, 1989) unless grown in 2i/lif condition(Morgani et al., 2013), it is unlikely that the T+ population represents an ExE-like population. However, to exclude this possibility we tested if T colocalised with Oct4 which is not expressed by the ExE (Downs, 2008). Our result shows that T+ cells found in Lif/FCS culture were also Oct4+ and the quantification revealed that the number of cells expressing the proteins correlated with the RNA expression profiles. These results suggest that the T+ population is likely to represent a PS progenitor population. We also found the Ki67 antigen to be expressed in T+ cells indicating that these cells are still actively proliferating (Fig. S1).

### Geometry dictates T patterning in ESC colonies

In vivo, the morphology of the embryo provides spatial constraints to shape morphogen gradients and to guide morphogenetic processes. Our previous results show that anterior and posterior genes are reciprocally regulated by diffusible signaling molecules. We also observed that a small proportion of cells expressing T at the protein level could be found in the culture (Fig. S1). However, in the dish, spatial disorganisation was evident. We hypothesized that the apparent spatial randomness observed in the dish was a consequence of the lack of geometric confinement. To test this idea, we developed a method to determine the preferential 3 dimensional distribution of cells in ESC colonies (Fig. S2). Briefly, we used micropatterns as a mean to provide geometrical constraints on the colonies. Micropatterns allow to precisely define the shape and the size of the area on which cells can adhere and grow. As a result, it becomes possible to project the localisation of the cells identified across multiple colonies onto one single shape. The result of this projection enables the generation of a probability density map (PDM) which corresponds to the preferential localisation of the cells within colonies.

We tested the effect of three distinct geometries onto the localisation of T+ cells. We designed shapes in order to conserve a constant area of 90 000 um2, while progressively introducing asymmetries in the geometry. When cells were grown on discs, the PDM clearly indicated that T+ cells were preferentially located at the periphery of the group after 48h of culture on patterns (Fig. 3). It is important to take into consideration that the PDM is a result obtained from accumulating the data from multiple colonies. We rarely observed an entire ring of T+ cells within one colony. Rather, we found small clusters of T+ cells which localised to the periphery, a result in agreement with a recent report (Tewary et al., 2017). To illustrate this, a representative z-stack is shown adjacent to the PDM in Fig. 3A. The mouse embryo possesses an ellipsoidal shape where the major axis aligns with the AP axis at the onset of gastrulation (Mesnard et al., 2004). We thus decided to test the effect on T localisation of a 2D ellipsoidal shape which includes similar changes in the curvature at the periphery. The PDM indicated that T+ cells could be found at the tips of the ellipse. This outcome was extremely reproducible and the PDM closely represented what could be observed by eye across entire fields of colonies within the dish as illustrated by the representative stack on Fig. 3A. Interestingly, some colonies contained T+ cells located only on one of the two sides of the ellipse. To determine if such a polarised expression of T could be obtained in a reproducible manner, we grew the cells onto half-ellipses as we reasoned that asymmetry would be reflected into the localisation of T+ cells. Indeed the PDM obtained on half ellipses demonstrated that T+ cells were localised on the tip of the half-ellipse. We did observe in some occasions that a few T+ cells were present in the corners of the shape, indicating that T+ cells were « attracted » by the regions with high curvature.

**Fig. 3.**
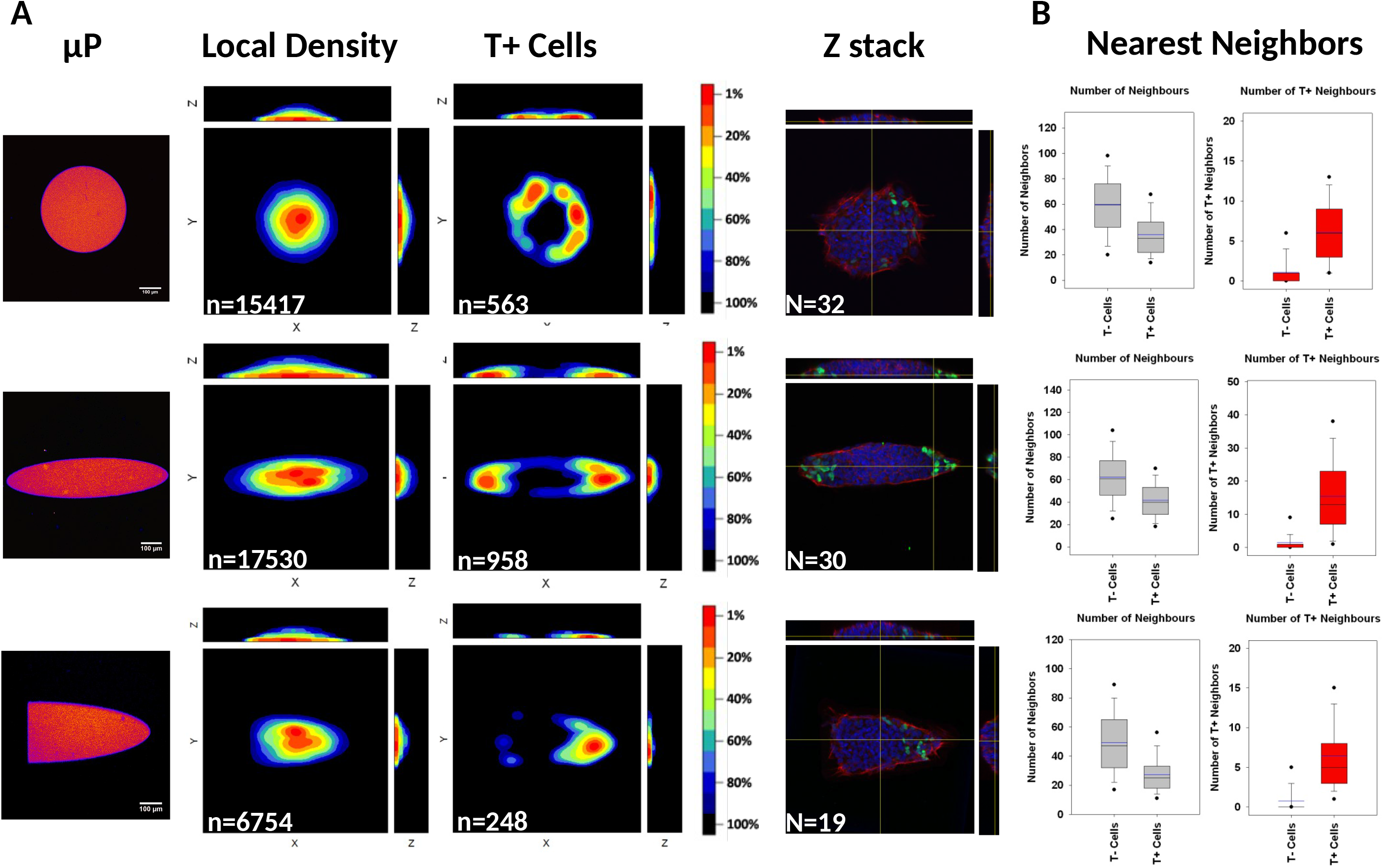
Geometry dictates T+ cells localisation in ESC colonies. Left column: confocal image of the micropattern’s autofluorescence showing the shape used for each row in the figure. Scale bar: 100µm. 2^nd^ column: PDM for every cell found in the samples. 3^rd^ column: PDM of T+ cells. The color table used for PDMs is represented, the scale indicates the percentage of cells found in the area depicted with the corresponding colour. 4^th^ column: representative z-stack image (blue: nuclei, red: Actin, Green: T). The last column shows the nearest neighbors analysis. For each cell, the number of other cell centroid comprised within a 30µm radius sphere was computed. The obtained distribution is represented as a Tukey box plot with symbols representing the 5 ^th^ and 95^th^ percentile. The black midle line is the median of the distribution and the blue line is the mean. The grey box plots represent the total number of neighbors for T- or T+ cells. The red box plots represent the number of T+ cells for T- or T+ cells. n = number of cells in the KDE analysis. N = number of colonies used for analysis.

Together our results demonstrate that spatial order and more specifically that the polarised positioning of T+ cells can be engineered by providing geometrical constraints to ESC colonies.

The fact that patterning occurs in ESC colonies on micropatterns proves that certain important features of the microenvironment are dictated by the shape of the colony.

Local cell density is amongst the candidate features which can regulate the positioning of T+ cells. Fig. 3A contains the PDM of the entire population of cells. By comparing the PDM of every cells versus the PDM of T+ cells, we could observe that the positioning T+ cells negatively correlated with local cell density on every shape. This was further confirmed by computing the number of neighbours found within a radius of 30 µm around the centroïd of each cell. Indeed, this quantification showed that regardless of the colony geometry, T+ cells had substantially fewer neighbours than T-cells. We also computed the number of T+ neighbours and found that T+ cells had more T+ neighbours than negative cells indicating that T+ cells are often found in clusters (Fig. 3B).

Another parameter which can be controlled by micropatterns is the repartition of forces across the colony. We observed large supracellular actin bundles lining the border of the micropattern (Fig S3), indicating that a regulated mechanical continuum was beeing established across the entire colony. We could also observe cell protrusions aligned with the direction of putative forces experienced by the cells of the periphery. Numerical simulations have suggested that cells experience high tension where the µP convex curvature is the highest (Nelson et al., 2005), which corresponds to regions where spread cells form large protrusions and where T+ cells localise.

### Endogeneous Nodal/Activin signaling regulates T+ cell number but not their spatial distribution

We have shown that paracrine signals regulate T expression in ESC culture and that T+ cells preferentially localise to regions of low density. This rises the possibility that morphogen gradients within the colony provide both differentiation signals and positional information leading to the observed distribution of T+ cells. As mentioned earlier, we found Lefty1 to be upregulated on micropatterns which could indicate an over-activation of the Activin/Nodal signaling pathway on micropatterns. In agreement, we found that Nodal transcripts were upregulated both at low density and on micropatterns (Fig. S4). Since the Nodal/Activin pathway has been shown to regulate T in ESC (Gadue et al., 2006) and since Nodal is required *in vivo* to pattern the pre-gastrulating embryo (Brennan et al., 2001), we decided to test the effect of perturbations of this pathway on the gene expression of ESC grown on micropatterns (Fig. 4A).

**Fig. 4.**
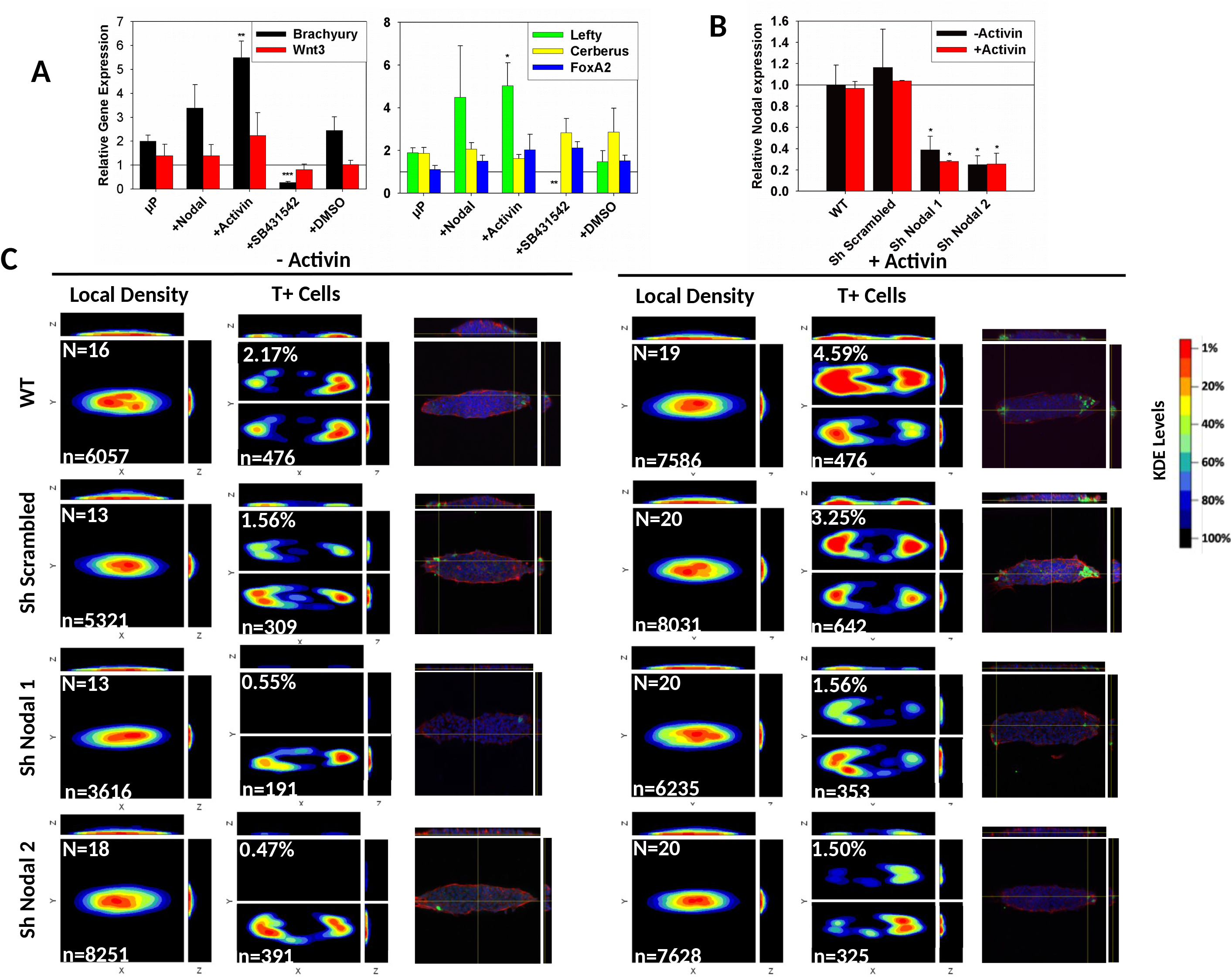
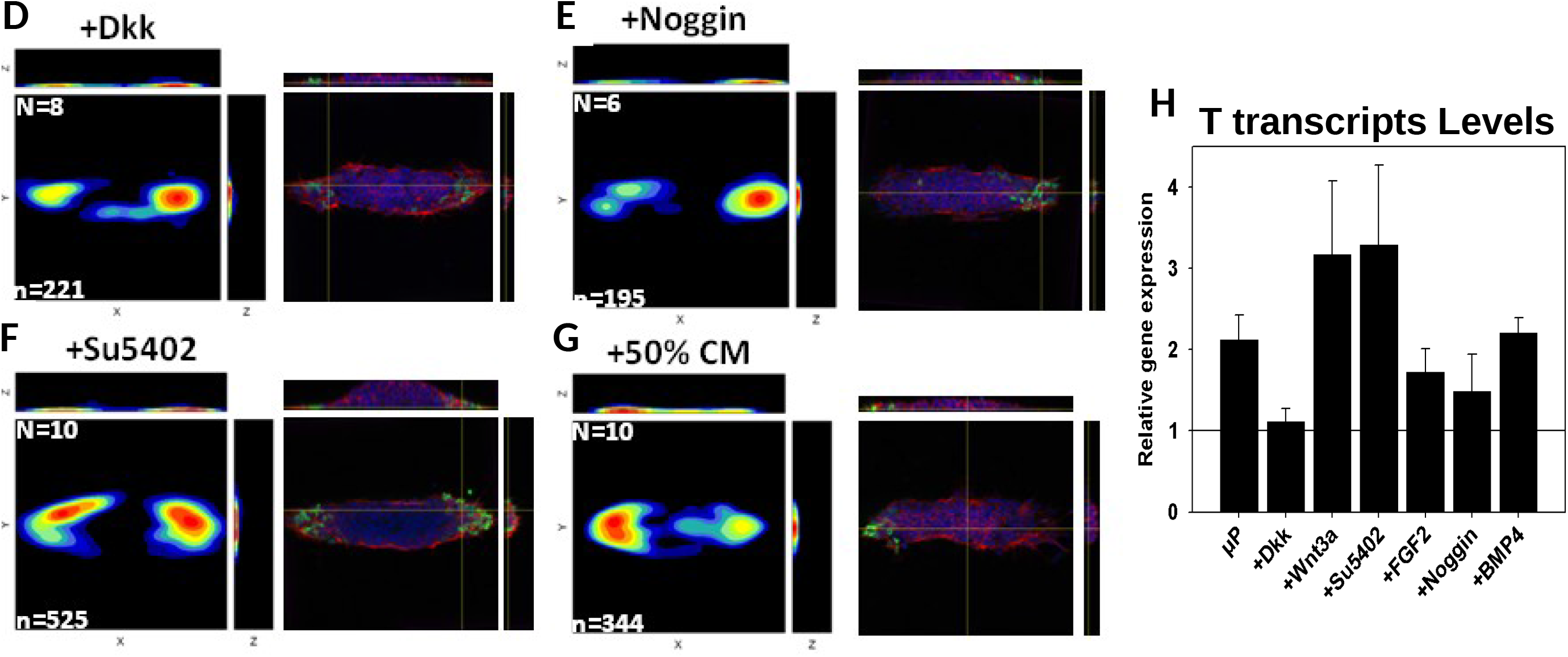
Paracrine signaling molecules modulate the number of T+ cell but not their spatial distribution. **A** qPCR analysis of proximal (left panel) and distal (right panel) gene expression in ESC grown in LIF/FCS for 48h on disc µP and treated with the indicated substances. **B** qPCR analysis of Nodal gene expression in WT cells or stable cell lines containing a non targeting shRNA or a Nodal targeting shRNA (2 distinct sequences were used). Error bars represent the S.E.M of at least 3 independent experiments. The stars represent the p value of the Student’s t-test as compared to low density (* p < 0.05, ** p < 0.01, ***p < 0.005). **C** WT cells or cell lines stably expressing shRNAs were grown for 48h on ellipsoidal µP with or without addition of recombinant Activin in the medium. For each condition, three panels are shown. Left: PDM for all the cells indicating local density. Middle: PDMs of T+ cells only Right: representative confocal image (blue: nuclei, red: Actin, Green: T). The middle panel is split into two sections, the bottom part shows an autoscaled PDM and the top section shows the normalised PDM values relative to the percentage of T+ cells found in WT without Activin: PDM level of sample = PDM level of sample x percentage of T+ cells in sample / percentage of cells in WT without Activin. n = number of cells used for KDE. N = number of colonies used in the analysis. The percentage of T+ cells in the sample is shown in the upper left corner of the T+ cells PDM. **D-G** Cells were cultured for 48h on ellipsoidal µP in the presence of the indicated inhibitor or in the presence of a mixture of fresh medium with conditionned medium (CM). **H** Levels of T transcripts relative to ATP50 in WT cells after 48h culture on ellipsoidal µP with or without treatment. The horizontal line indicates the level of T expression in the unpatterned control condition. Error bars represent the S.E.M of at least 3 independent experiments.

We observed that addition of either 1µg/ml of recombinant Nodal or 25ng/ml of Activin A increased markedly the level of expression of both T and Lefty1. Conversely adding 10µM of SB431542, a potent inhibitor of Nodal/Activin signalling, strongly reduced the level of T and totally abrogated Lefty1 expression consistently with the fact that Lefty1 is a direct target of the Nodal/Activin pathway. In contrast, we did not observe any significant effect of recombinant Activin/Nodal or SB431542 on the levels of expression of Wnt3, Cerberus or FoxA2 indicating that these genes are regulated by other factors. We noticed that a much higher concentration of recombinant nodal was required to induce an effect. However, this is consistent with a work based on a smad2 responsive luciferase reporter showing that Activin is about 250 times more potent than Nodal to activate the pathway(Kelber et al., 2008).

Motivated by the above observations, we decided to investigate whether an endogenous gradient of Nodal signaling could be involved in the regulation of T+ cell number and distribution. To address this point, we generated stable cell lines knocked down for Nodal by RNA interference. We confirmed the decrease of Nodal transcripts in the cells knocked-down for Nodal (Nodal-KD) compared to the level of WT cells or to the level of a cell line carrying a non targeting shRNA construct (Fig. 4B).

To monitor the effect of Nodal knock-down in ESC, we seeded cells on ellipsoidal µP and constructed the PDM for T+ cells after 48h (Fig. 4C). We observed a 3 to 4 fold reduction in the number of T+ cells in Nodal-KD cells. In order to create a visualisation which accurately represents such changes across conditions, we normalised the levels of the PDM to the levels observed in the WT condition without Activin. Auto-scaled PDMs are also provided to allow the visualisation of the preferential localisation of T+ cells when very low number of T+ cells were observed (Fig. 4C). Remarkably, although the number of T+ cells was lower in Nodal-KD cells, the positioning of T+ cells remained unaltered. This observation might still be explained by a gradient of low Nodal activity across the colony which is sensed by cells with a low threshold response. To attempt to override the putative gradient of Nodal activity in the colony, we treated the cells with Activin A. Importantly, unlike Nodal, Activin A is not affected by Lefty1 inhibition (Chen and Shen, 2004). We found that Activin A treatment increased the number of T+ cells in WT or control cells and that it rescued original T+ cell number in Nodal-KD cells. Most importantly, the positioning of T+ cells was not affected. These results demonstrate that Nodal signaling can regulate the number of T+ cells in the colonies but it does not dictate their positioning.

We further interrogated the role of other signaling molecules which could potentially be defining the positioning of T+ cells, including Fgf, Wnt and BMP. We first tested the effect of treating the cells with either agonist or antagonist of each pathway onto gene expression. We observed a reproducible increase of T transcripts upon treatment with Wnt3a, which was confirmed by the downregulation of T when the cells were treated with Dkk (Wnt inhibitor). This corroborated the idea that a low level of endogenous Wnt participates in the regulation of T expression. Interestingly, we found that inhibition of Fgf signaling with SU5402 also slightly increased the number of T transcripts, indicating that endogeneous Fgf may be a negative regulator of the number of T+ cells. On the other hand, we did not observe any significant effect of either exogeneous Fgf, BMP or Noggin (BMP inhibitor). This may be attributed to the failure of these recombinant proteins to further modulate an already strongly active endogenous level of signaling.

We then treated cells grown on micropatterns with either SU5402 (Fgf inhibitor), Dkk1 (Wnt inhibitor) or Noggin (BMP inhibitor) and created PDM for T+ cells (Fig. 4 D-F).None of the pathway inhibitors impacted the localisation of T+ cells indicating that neither, Fgf, Wnt or BMP were responsible for T patterning on micropatterns. We finally tested the effect of CM and found no significant evidence of patterning disruption. These observations tend to suggest that localisation of T+ cells on the other hand, is not determined by self-generated gradients of morphogens in the colony.

### PS progenitors patterning involves cell movement and not a change in gene expression

Patterning of T+ cells on micropatterns may be established either via de novo T expression at the tips of the ellipse or via positional reorgnisation of pre-existing T+ cells. To discriminate between these two hypotheses, we manipulated the initial proportion of T+ cells seeded on µP. We reasonned that if cells were turning T on and off, the proportion of cells found at the end of the experiment should be independent of the initial proportion of T+ cells in the starting population. Conversely, if T+ cells movement was responsible, the proportion of T+ cells should remain approximately constant and cells should be found at the tips of the shape (Fig. 5A). We showed previously that the amount of T+ cells within the dish could be increased or decreased significantly by culturing cells at low or high density respectively (Fig. 2C). Thus, we first tested if the proportion of T+ cells in the population was reversible when cells were switched from one density to the other (Fig. 5B). We found that the proportion of T+ cells grown at low density and then at high density changed from 25.1% to 2.3% whereas the proportion of T+ cells grown first at high density changed from 0.6% to 12.5%. This result demonstrates that preconditioning the cells does not alter their capacity to respond to density and that the proportions of T+ cells can rapidly change in response to environmental changes. Therefore we tested next the effect of preconditioning cells at low or high density prior seeding onto µP. We observed that 19.5% of the cells were T+ when cells came from low density culture whereas only 1.7% were T+ when cells came from high density culture. Most importantly, regardless of the initial conditions, T+ cells were excluded from the colony center (Fig. 5B). This result shows that only a minor proportion of cells are turning T on or off when they are transferred on µP and that the majority of T+ cells initially present at the start of the experiment are still present at the end. This observation is also in line with the fact that we did not observe any negative correlation between the total number of cells and the number of T+ cells in the colony (Fig. 5C), a phenomenon which should be observed if gradients of morphogen activity generated at the colony level were sufficiently potent to switch the fate of the majority of the cells. Thus, this experiment is strongly in favour of a mechanism involving cell repositioning.

**Fig. 5.**
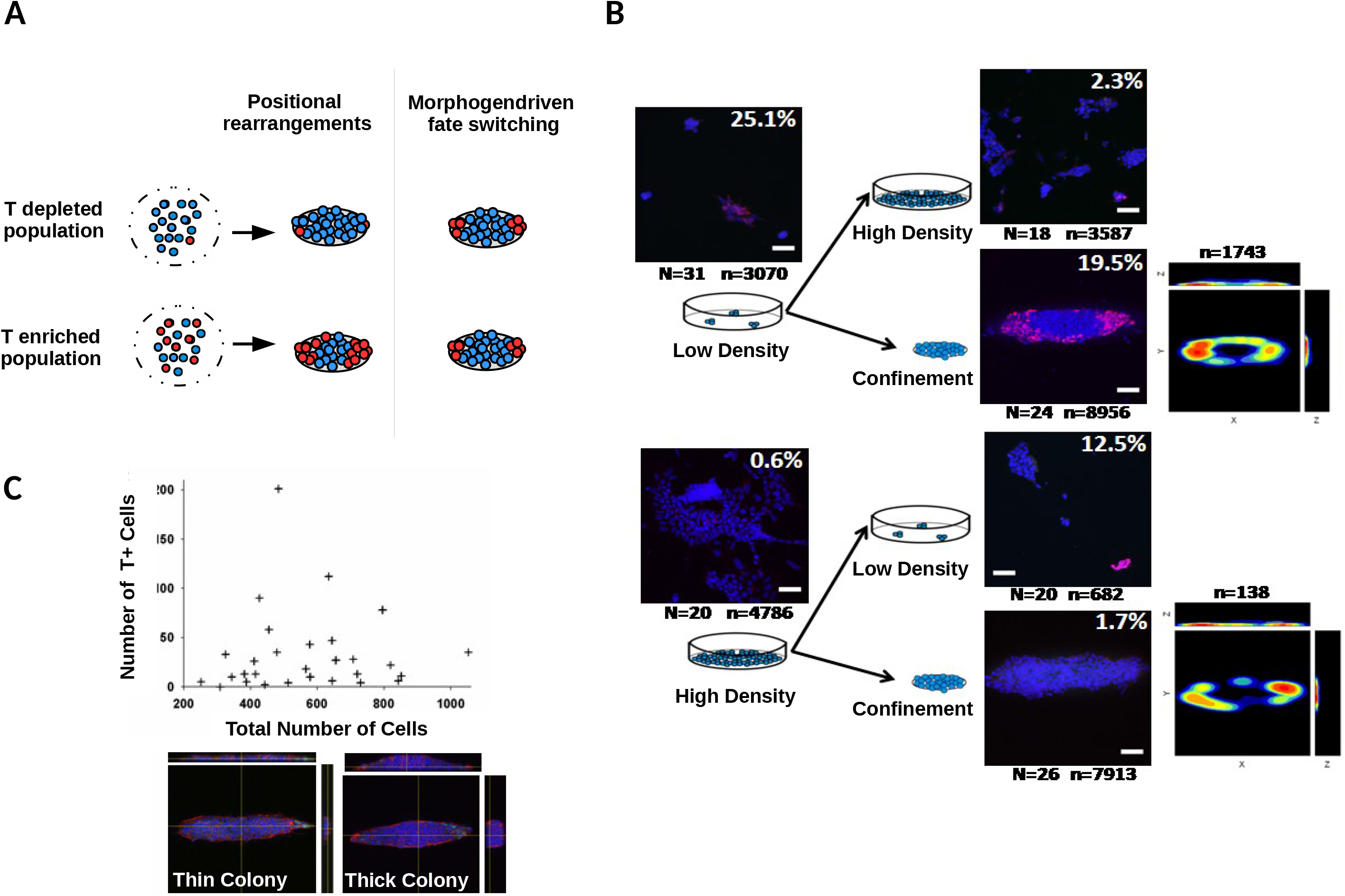
PS progenitors patterning involves cell movement and not a change in gene expression. **A** Scematic illustrating the rationale to distinguish between two possible mechanisms involved in the patterning of T+ cells. If self-organised gradients of morphogen switch the fate of the cells in a position dependent manner, the final number of T+ cells should not depend on the initial number of T+ cells. If cell movements instead are involved, the T+ cell number should remain constant. **B** Cells were grown at low or high density in order to obtain a population of cells that is enriched or depleted of T+ cells respectively. Each sample was then replated either on µP or at a reciprocal cell density. Analysis for T+ cell content and patterning was performed after 48h. The percentage of T+ cell in the population is indicated in the upper left corner of the representative confocal images (blue:dapi, red: T), scale bar: 50µm. **C.** Number of T+ cells plotted against the total number of cells within the colony where each data point represent one colony. Two representative colonies are shown on tunderneath the graph. Note the difference in thickness indicating the difference in the number of cells in each colony.

### The positioning of T+ cells is a robust and rapid process

To obtain some insights about the kinetics and the robustness of T+ cells patterning, we randomised the position of T+ cells from patterned colonies and performed a time course to monitor the reestablishment of patterning over time (Fig. 6A). After 4h we could observe T+ cells embedded within clumps of negative cells showing that T+ cells were not lost during the process. Next at 24h, we could see that T+ cells started to be excluded from the denser parts of the colony. Finally, at 48h, T+ cells were found only at the tips of the ellipse. This experiment demonstrates that the positioning of T+ cells within geometrically confined ESC colonies is very robust and that the exclusion from the dense center of the colony is a process which takes less than 24h. The localisation of T+ cells at the tips of the ellipse was then further refined between 24h and 48h.

**Fig. 6.**
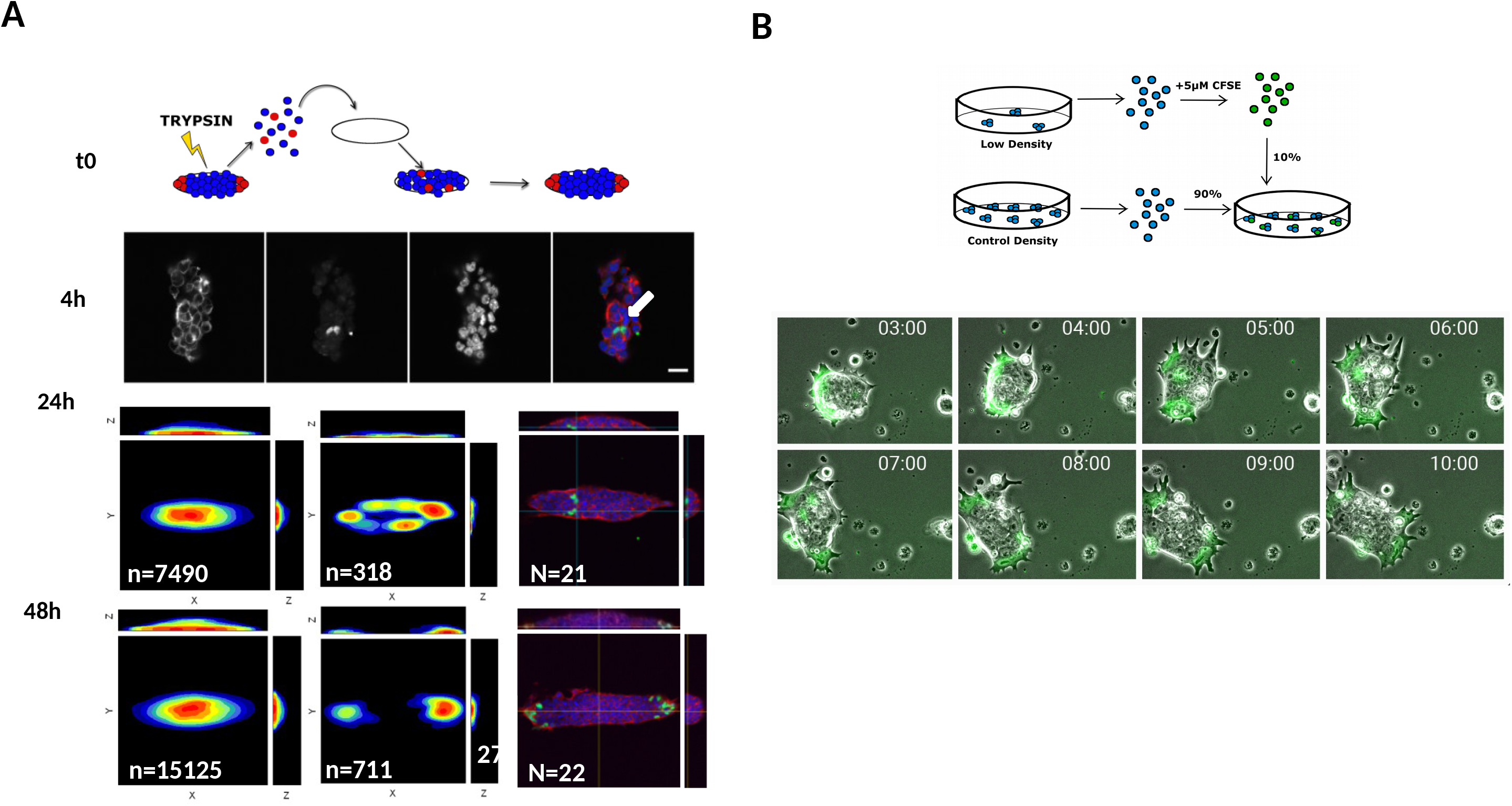
T+ cells patterning is a robust and rapid process. **A** Cells were cultured for 48h on ellipsoidal µP to allow patterning of T+ cells. Then, cells were trypsinized and seeded again on fresh µP in order to randomise the position of T+ cells within the group. Cells were fixed after 4h, 24h and 48h, stained for T and analysed. At 4h, the white arrow indicates T+ cells embedded within the group, scale bar: 20µm. At 24h and 48h, the left panel is the PDM for every cells, in the midle, the PDM for T+ cells and the right panel is a confocal image of a representative patterned colony. **B** Cells were grown at low density in order to obtain a population enriched in T+ cells. This population was then stained with CFSE and mixed with unstained cells in a 10:90 ratio. The cells were then allowed to adhere and were imaged 4h after plating every 3 minutes. The sequence of images illustrates the rapid positioning of the T enriched population at the periphery of the colony and shows how this population guides the movements of the colony (see also Supplementary movie 2).

Importantly, we noticed that contrary to the 48h situation, a large proportion of patterns were not fully covered by the cells at 24h even though the patterns were almost entirely covered by cells a few hours after plating. We explain this by the fact that ESC express high levels of E-Cadherin and therefore tend to form tightly packed colonies. Sometimes, two groups of cells could form on one pattern and then merge as they grow as shown in sup movie 1. This effect may bias our interpretation of the real kinetics of the exclusion of T+ cells from the center of the colony as cells localised at the periphery of one colony could become trapped in between two merging colonies at around 24h. In order to observe a more direct evidence of cells reorganisation, we performed a time lapse experiment where we plated a mixture of CFSE stained cells previously grown at low density (to obtain a T+ enriched population of cells) together with unstained cells previously grown at a control density (Fig. 6B). This experiment showed striking evidence that some CFSE+ positive cells were very motile, projecting protrusions at the periphery of the colony. These cells seemed to undertake the role of guiding the migration of the entire colony. Importantly, we observed that some CFSE+ cells could elongate to “extrude” themselves from dense regions of the colony (Sup Movie 2).

Together these data provide strong evidence that the positioning of T+ cells on micropatterns is a result of cell movements and is not due to de novo T expression in situ. This suggests that T+ cells have acquired the competence to sense the surrounding of their environment and interpret geometrical cues.

## Discussion

Between E5.5 and E6.5 of mouse development, AP polarity develops from a proximo-distal (PD) pattern of gene expression. T and Wnt3 transcripts are detected posteriorly whereas Lefty and Cerberus transcripts are expressed at the distal tip of the embryo. This PD pattern of gene expression is established by the combined actions of inductive signals and negative feedback loops which generate two domains that mutually repress the expansion of the other (Arnold and Robertson, 2009; Rossant and Tam, 2009).

Here we provide evidence that the reciprocal regulation of proximal and distal genes is recapitulated within ESC cultures, enabling anterior and posterior cell types emerge in particular proportions. We discuss how the emergence of signalling networks might control this behaviour in section 1 of the discussion

However, whilst signalling networks explain the proportions of different cell types, they do not explain their spatial organisation, rather this may be influenced by cell geometry guilding cell movements. We discuss possible mechanisms underlying this behaviour in section 2 of the discussion.

### Regulation of PS and DVE identities in ESC cultures

By culturing ESC at different densities in Lif/FCS condition, we evidenced that the reciprocal regulation of proximal and distal genes is recapitulated within ESC cultures. This poses the question of how this balance is regulated in LIF/FCS ESC culture.

### Induction of posterior markers

In the absence of signals, ESC differentiate towards the anterior neural fate (Hemmati-Brivanlou and Melton, 1997) and lif alone does not sustain pluripotency. Thus serum contains an agent which antagonise the neural lineage. The main candidate is proposed to be BMP since BMP can substitute for serum to maintain ESC in culture with Lif (Morikawa et al., 2016; Ying et al., 2003b). In vivo, BMP elicit Wnt3 and sustains Nodal expression to confer a proximo-posterior identity to the epiblast (Ben-Haim et al., 2006). We observed that T which is a known target of the canonical Wnt pathway (Arnold et al., 2000) was indeed dependent on endogeneous levels of Wnt in our culture. We also confirmed that manipulating Nodal using recombinant proteins, inhibitors or shRNA could modulate the level of T expression. Finally, we observed that culturing the cells at low density increased the amount of Nodal and Wnt 3 expressed by the cells. These data are all in agreement with previous reports(ten Berge et al., 2008; Turner et al., 2014b) that the BMP/Nodal/Wnt autoregulatory loop is recapitulated in ESC.

### Upregulation of anterior markers and downregulation of posterior markers at high density

At high density on the other end, T and Wnt3 were downregulated in favour of the VE marker Sox17 and the AVE markers FoxA2 (Kimura-Yoshida et al., 2007) and Cerberus (Belo et al., 1997). We also report the detection of a Cerberus+ population of cells in EBs and FACS followed by qPCR analysis shows that this population is enriched in FoxA2 transcripts (Fig. S7).

We would like to speculate that the detection of these markers indicates the presence in the culture of a small subpopulation of cells acquiring organiser activity. Although further experiment will be required to definitively demonstrate this point, this hypothesis is supported by the work from other groups who have reported independently the promoter activity of VE genes, Hex1(Canham et al., 2010) and Sox17 (Niakan et al., 2010) within small fractions of ESC. The thorough characterisation of these subpopulations have shown in both cases that these cells were primed towards the extraembryonic endoderm lineage as they were able to efficiently contribute to the VE lineage and its derivatives in Chimeras.

With this in mind, we propose that the mechanisms that lead to the expression of AVE markers in high density culture involves a sequence of events which relatively closely resembles situations happening in vivo (Fig. 7). Indeed, during normal propagation, ESC experience diverse microenvironments as cell densities are not homogeneous within the dish (Davey and Zandstra, 2006). In addition, some degree of stochastic gene expression participates in the generation of a range of cell competence to respond to signaling as seen in vivo (Chazaud and Rossant, 2006; Grabarek et al., 2012; Hermitte and Chazaud, 2014; Yamanaka et al., 2010). This translates into the presence in the dish of a PE specified subpopulations of cells. The proportion of these cells is likely to be regulated by Fgf signaling. Fgf is the main pro-differentiation signal secreted by ESC (Kunath et al., 2007), and is required for the derivation of extraembryonic endodermal cells from ESC (Cho et al., 2012; Niakan et al., 2010). In addition, Canham and colleagues have demonstrated that Fgf signaling controls the size of the Hex+ subpopulation (Canham et al., 2010) and in vivo Fgf specifies the PE in the ICM (Chazaud and Rossant, 2006; Grabarek et al., 2012; Hermitte and Chazaud, 2014; Yamanaka et al., 2010).

**Fig. 7.**
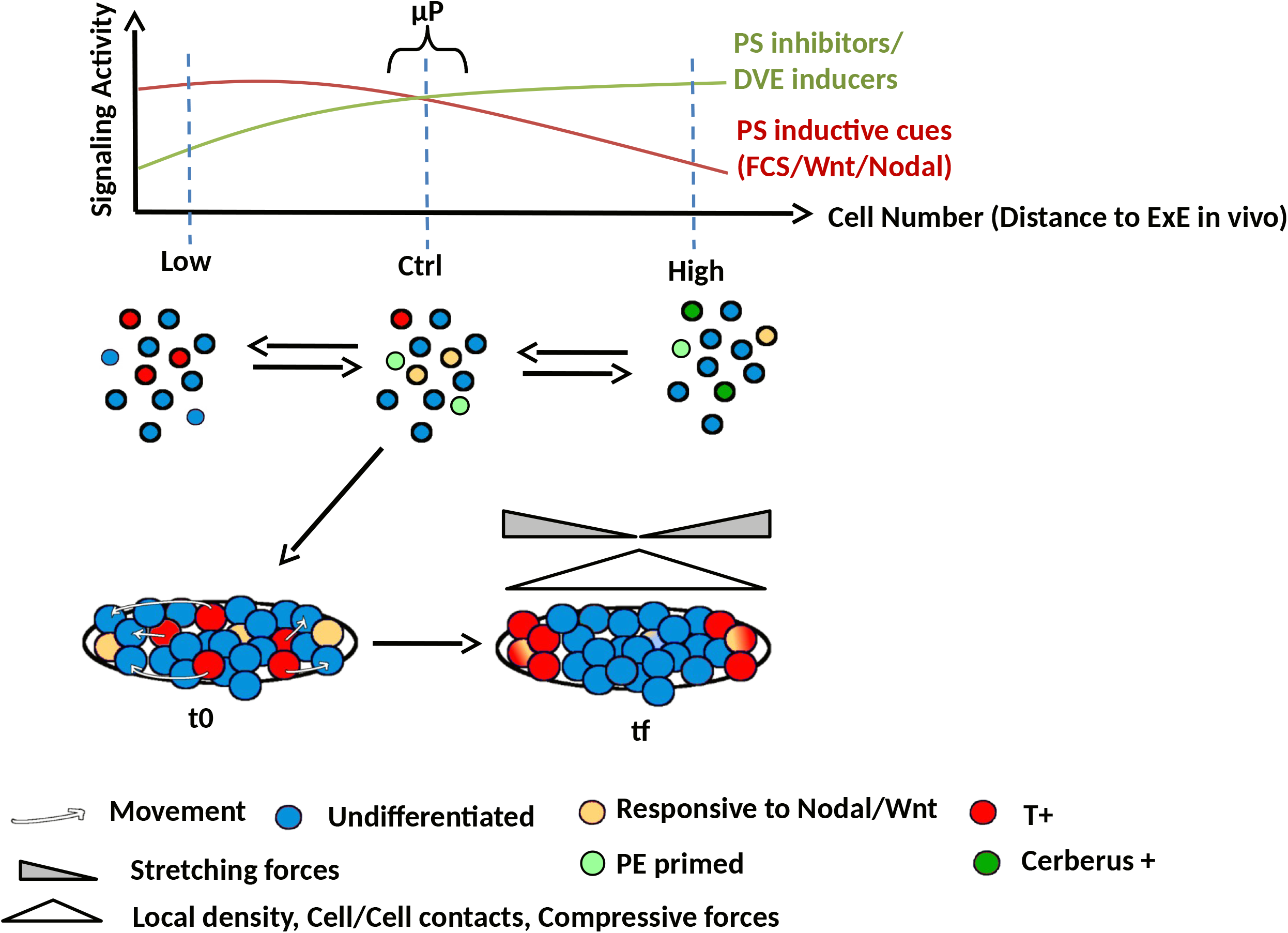
Model of PS and AVE gene regulation in ESC and T+ cells patterning with confinement. ESC in culture consist in a mixture of discrete cell states where distinct subpopulations of cells possess various level of competence to respond to the cues from their environment. The proportion of each subpopulation within the culture depends on the biochemical signals provided by the medium and the cells themselves. The upper graphic illustrates how ESC culture density modulates signaling activity and the schematics below how the proportion of several identified and putative subpopulations are affected. Importantly, the signals and the resulting cell states are developmentally relevant. During normal propagation of ESC, FCS prevents the differentiation of cells towards the neural lineage while promoting a more proximoposterior identity to the cells. At an optimal cell density, Lif and possibly juxtacrine gp130 signaling prevent differentiation to mesendoderm while signals from the cells also damper FCS signals. At high density on the other end, FCS and Lif signaling become limiting. FCS antagonising factors secreted by the cells become sufficient to promote a more anterior identity to the cells. On µP, the specific cellular arrangement of the cells tend to potentiate both signaling activities. An effect which in lif/serum condition is not sufficient to have a significant impact on cell identity. When ESC cells are provided with geometrical cues on micropatterns, T+ cells are able to sense the geometry and forces of their microenvironment and migrate towards the low density regions of the colony, where stretching forces are the highest. A minor contribution of cell identity change is also represented. t0: time at the beginning of the experiment, tf: time at the end of the experiment.

Further specification of the PE into VE and then into Cerberus or FoxA2 expressing cells requires that the cells are exposed to additional signals. The secretion of laminin by the cells themselves appear to play a significant role (Paca et al., 2012). In vivo the specification of the VE into DVE cells requires Nodal while BMP from the ExE restricts its formation to the distal tip of the embryo (Brennan et al., 2001; Clements et al., 2011; Rodriguez et al., 2005). The picture is however complexified by the fact that cells which participate to the AVE have multiple origins and Cerberus expression for example can be tracked back to the pre-implantation stage (Granier et al., 2011; Takaoka et al., 2011; Torres-Padilla et al., 2007). Importantly, DVE cells may be specified either via stochastic signaling threshold reaching events involving Nodal and Lefty at the blastocyst stage or via a drop of BMP signaling activity away from the source at the peri implantation stage (Hiramatsu et al., 2013; Stower and Srinivas, 2014).

Thus, in the culture, as Nodal is actively driven by the pluripotency GRN (Papanayotou et al., 2014), paracrine Nodal can further induce PE biased cells to acquire VE characteristics. In this model, BMP from the serum and Lif prevent the overt differentiation to DVE in normal condition. However in the HD situation, one can imagine that the supply in Lif and BMP becomes limiting. Indeed, several clues indicate a BMP inhibition at high density. First, both Cerberus and Sox1 expression require an inhibition of the BMP pathway (Malaguti et al., 2013; Soares et al., 2008; Ying et al., 2003b). Second, the amount of Nodal and Wnt3 transcripts correlated negatively with density suggesting that the signal maintaining the Nodal/Wnt autoregulatory loop at HD was being deconstructed. And finally, CM from high density to low density culture switched off the expression of posterior markers in favor of an increase in expression of Cerberus. The exact paracrine signals responsible for this effect remain to be determined. A BMP induced negative feedback loop has been demonstrated in human ESC (Etoc et al., 2016; Tewary et al., 2017; Warmflash et al., 2014) which nottably involves Noggin. Interestingly mouse EpiSC did not secrete Noggin in response to BMP4 (Etoc et al., 2016), however other BMP inhibitors such as GDF3 which are expresed in mESC (Levine and Brivanlou, 2006) may participate.

In the embryo, Lefty1 and Cerberus mark distinct populations of the DVE (Stower and Srinivas, 2014; Torres-Padilla et al., 2007) and indeed, Lefty1 did not follow the pattern of expression of Cerberus in our system either. In ESC however, Lefty1 is both associated with stemness and upregulated during differentiation (reviewed in (Tabibzadeh and Hemmati-Brivanlou, 2006)), thus the use of Lefty1 as a marker of the DVE in this context remains questionnable. Lefty1 is also a direct target of the smad2/3 pathway (Besser, 2004), and we indeed observed a very strong effect of Nodal modulators on the level of Lefty1 transcript. The fact that Lefty1 does not change with CM indicates that Nodal activity is not directly affected by paracrine signals. This rules out Nodal inhibitor such as Lefty1 itself or Cerberus which is also known as a Nodal Regulator (Perea-Gomez et al., 2002) as the main drivers of the effect of CM.

Another possibility is that CM contains inhibitors of the canonical wnt pathway. CM did not impact regulation of Lefty1 while it impacted T regulation, similar to our treatment with Dkk1. It will be interesting to test the amount of Dkk1 expressed by the cells at HD since DVE cells antagonize posterior fate by secreting Dkk1 *in vivo(Glinka et al., 1998)*.

Finally, although we showed that diffusible molecules are the main driving force in ESC culture as evidenced by CM experiments, cell response to posteriorising signals may as well be affected by YAP/TAZ activity and receptor presentation in HD cultures as reported for other systems (Azzolin et al., 2014; Dupont et al., 2011; Etoc et al., 2016; Narimatsu et al., 2015; Varelas et al., 2010; Warmflash et al., 2014).

Altogether our data provide a novel evidence for the idea that ESC encompass discrete cells states representative of various developmental stages, including extraembryonic lineages, and that the molecular machinery which defines the AP polarity in the embryo is also at work in conventional ESC culture.

These considerations may have important implications for the understanding of the self-organising ability of ESC organoids. Indeed, due to the observation that ESC only participate to the epiblast in chimera, the ability of ESC to generate extraembryonic lineages has been somewhat under-appreciated. It would be interesting to observe the behaviour of a Cerberus reporter in gastruloids(Brink et al., 2014) or in TSC/ESC aggregates(Harrison et al., 2017). Indeed, Sox17 which marks both the VE and the definitive endoderm (Ref Clements) was found to be polarised in mouse Gastruloids. Reconsideration of the fact that extraembryonic endoderm can emerge from ESC may clarify how spontaneous symmetry breaking events can occur in ESC derived organoids in the absence of extraembryonic lineages of embryonic origin.

### Mechanisms of T+ cells patterning

When ESC were deposited on uP, we observed a 2 fold increase of both PS markers and anterior markers compared to the control condition. Interestingly, Lefty1 expression was also higher on µP than in any other condition, possibly indicating a slight over-activation of the Nodal signaling pathway. The confinement of the cells on micropatterns did force the cells to aggregate and to form dense colonies with multiple layers of cells. At the same time, the overall cell number in the dish was similar to the control condition, thus avoiding a global depletion of exogeneous factors. Given the considerations discussed earlier, we may wonder if the spatial confinement of ESC on uP was sufficient to create the distinct cellular microenvironments required for both PS and AVE specification trajectories to occur within one single colony. We believe however, that in our conditions the cellular specification to either fate due to these microenvironmental differences is marginal. One of the main evidence for this is that when cells were preconditionned at low or high density, only extreme changes in the total number of cells in the dish could dictate the number of T+ cells, whereas on patterns the rate of T expression reversal was very small in comparision.

This emphasizes the major differences which distinguish the present work from the **previous** studies plublished recently on peri-gastrulation like events on micropatterns with human ESC. In these articles, unspecified hESC are released from their pluripotency states by removal of Fgf and Activin, thus, the process investigated involves the emergence of cell fates in response to self organised gradients of signaling in response to BMP4. Long range gradients are allowed to form thanks to the size of the patterns (∼1mm) and indeed Tewary and colleagues have confirmed that patterns could scale with larger patterns but not with smaller ones (such as ours). Therefore, these reports describe a faithful and quantifyable model of germ layer patterning in vitro which allow to interrogate the mechanisms of gradient formation and cell response to these gradients.

In the present study, we ask a distinct question, which is the influence of tissue geometry in guiding self-patterning when multiple cell states that have already been specified by morphogens are present in a spatially disorganised population. Indeed, we have systematically tested the roles of diffusible signals in this process and shown that although signaling molecules were regulating the number of T+ cells, none of the disrupted pathways had a significant impact on T+ cells localisation (Fig. 4). These observations lead to the conclusion that cell movements are responsible for the patterning of T+ cells on uP. Multiple mechanisms can account for positional reorganisation. Cell sorting can be dictated by differential cell to cell cohesion and cell contractility (Foty and Steinberg, 2005; Krieg et al., 2008; Lecuit, 2008). ESC possess a high level of E-Cadherin while PS cells undergo an EMT which in the streak involves a switch from E-Cadherin to N-Cadherin (Radice et al., 1997) and an increase in cell contractility (Burute et al., 2017; Tseng et al., 2012). Aditionally, Nodal has been shown to regulate cytoskeletal reorganisation and dynamic (Krieg et al., 2008; Trichas et al., 2011). Thus, differential cortical tension and inter-cellular adhesion may participate to the spatial segregation of T+ cells to the lowest cell density regions of the colony.

An alternative, although, not exclusive explanation is that T+ cells have acquired a competence to actively migrate and to be guided by confinement. T expression has long been associated with motility (Hashimoto et al., 1987; Yanagisawa et al., 1981). Indeed, cells mutant for T fail to robustly contribute to mesodermal lineages in chimeric experiments due to a defect in migrating away from the midline (Wilson et al., 1995). Importantly, T over-expression in epthelial cancer cells promotes EMT and migration (Fernando et al., 2010), and ChiP-seq analyses have shown that T-bound regions are enriched with the promoters of EMT and migration related genes both in human (Faial et al., 2015) and mouse ESC (Lolas et al., 2014). Thus, T is not only a marker of the PS but also a transcription factor which contributes in altering the adhesive and migratory behaviours of the cells.

Migration can be guided by morphogens. Fgfs molecules are chemotaxic cues which guide the movement of T+ cells after ingression (Sun et al., 1999; Yang et al., 2002). Since ESC secrete Fgf4 (Kunath et al., 2007) (which repel T+ cells in the chick), one could imagine that T+ cells are repulsed from the colony center where the cells are the denser. However, we excluded this possibility as treatment with the Fgf receptor inhibitor SU5402 did not perturb the localisation of T+ cells (SI6). Instead, T+ cells migration may be sensitive to the distribution of forces dictated by the shape of the colony. In support of this idea, we observed supracellular actin networks spanning the entire colony with large actin cables lining the pourtour of the pattern (SI 3). This architecture indicates the generation of a regulated mechanical continuum across the colony which is highly reminiscent of multicellular actin network observed in models of collective cell migration (Mayor and Etienne-Manneville, 2016). Consistently, our time lapse experiment shows how ESC migrate and explore their environement as a cohesive unit with T-enriched cells quickly localising at the tip of the colony possibly participating in guiding the movement of the group, as it has been shown for epithelial cells expressing high level of RhoA and exerting strong traction forces on the substrate (Reffay et al., 2014). The shape of the colony, by orienting the intercellular forces within the colony and the traction forces at its periphery can direct these contractile leaders cells to emerge from high curvature regions (Rolli et al., 2012).

Such a process of tissue geometry sensing may have important implication during the early stages of gastrulation. Some recent evidence may suggest that EMT (or polarity reversal) positively regulate the level of T expression (Burute et al., 2017; Turner et al., 2014a). As both mesoderm and endoderm emerge from T+ cells (Rodaway and Patient, 2001), an interesting question is whether the geometry sensed by PS cells which have ingressed in the streak can provide feedback onto downstream cell fate decisions as well as participating in the spatial segregation of both layers.

A final comment maybe raised regarding the final positioning of the AVE during the establishment of AP polarity in vivo. Indeed, the AVE requires active cell migration (Migeotte et al., 2010; Rakeman and Anderson, 2006; Srinivas et al., 2004; Stower and Srinivas, 2014; Trichas et al., 2011), however the cues which guide the direction of the migration are still unknown. It will be interesting in the future to investigate the positioning of VE reporter cells on asymmetric patterns to determine if our observation with T+ cells can be reproduced with another relevant cell type. Much efforts have been put in order to determine if extrinsic cues were necessary to guide AP polarity in vivo. Mesnard and colleagues have shown that AP polarity does not align with the uterus axis but rather aligns with the main axis of the ellipsoidal shape of the embryo (Mesnard et al., 2004). More recently, the stiffness of the tissue surrounding the embryo was proposed to be required for the establishment of the DVE (Hiramatsu et al., 2013). This view has however been challenged by the fact that DVE appearance and proper AP polarity emerges within cultured embryo in the absence of external physical constraints (Bedzhov et al., 2015) even though, absence of a stiff support appears to be an important requirement for proper development of embryos in vitro (Morris et al., 2012). These evidence demonstrate that AP axis formation is an embryo-autonomous process and indicate that shape of the embryo and physical constraints may play an important role. The possibility that external cues from the mother provide additional robustness to embryonic development should not be completely ruled out.

### Conclusion

In this study, we provided geometrical confinement to a spatially disorganised population of cells and found that a subset of this population expressing characteristic primitive streak genes acquired the competence to be guided by this confinement. This resulted in the robust localisation of these cells towards the region where high matrix/cell tension is expected. We believe these result may have important implications regarding how T+ cells behave in the streak and how robustness at the onset of gastrulation may be achieved. Taken together our results are in agreement with the idea that ESC are endowed with the capacity to both respond and generate signaling cascades which correspond to the machinery employed by the embryo around peri-implantation stages. An engineering perspective consists in interogating which constraints need to be provided to this system in order to reanimate the construction of an ordered and organised sequence of events which mimicks embryonic development. Together with recent findings and the possibility that AVE-like cells emerge from ESC cultures, this work contributes in shaping the path towards an in vitro model of AP polarity.

## Materials and Method

### ESC culture and plasmid

ES cells (CGR8) were propagated in BHK21 medium supplemented with pyruvate, non-essential amino acids, mercaptoethanol 100µM, 7.5% fetal calf serum and LIF conditioned medium obtained from pre-confluent LIF-D cells stably transfected with a plasmid encoding LIF. The cells were trypsinized and replated 1/6 every two days. Knocked down cell lines were established and routinely passaged at the exact same density and the same day as a control cell line transfected with the sh Scrambled plasmid. All shRNA constructs were created using the psiRNA-h7SKblasti G1 expression vector (Invivogen) according to the manufacturer’s protocol. Target sequences for short hairpin RNAs are as follow: shControl: GCATATGTGCGTACCTAGCAT (*scrambled* oligonucleotide sequence, available prepackaged in the *psiRNA*-h7SKblasti vector, Invivogen), shNodal 1: GAAGGCAACGCCGACATCATT, shNodal 2: GGGAGCAGAAACGTTAGAAGA.

### Real time PCR

RNA was extracted from ES cells using a Zymo research kit. One µg of RNA was reverse-transcribed using the SuperscriptII reverse transcriptase (Invitrogen, Cergy, France) and oligodT_(12-18)_. qPCR was performed using a Light Cycler LC 1.5. Amplification was carried out as recommended by the manufacturer. 12 µl reaction mixture contained 10 µl of Roche SYBR Green I mix respectively (including Taq DNA polymerase, reaction buffer, deoxynucleoside trisphosphate mix, SYBR Green I dye, and 3 mM MgCl2), 0.25 µM concentration of appropriate primer and 10 ng of diluted cDNA. Melting curves were used to determine the specificity of PCR products, which were confirmed using conventional gel electrophoresis and sequencing. Data were analysed according to Pfaffl using ATP50 as the reference gene.

### ESC Micropatterning

A non fouling surface was obtained by incubating 2cm X 2cm hydrophobic ibitreat slides (ibidi, France) with a solution of 0.4% pluronic acid in PBS (sigma, France) for 30 min. Then, to remove excess of pluronic acid, the slides were rinsed twice with PBS and air dried. In order to create hydrophilic patterns, the slides were illuminated with deep UV light (UVO Cleaner, Jelight, USA) for 5min through a custom photomask (Toppan Photomasks, France). Finally, micropatterned substrates were coated with 0.1% gelatin (sigma, France). To seed the cells on micropatterned substrates, the slides were put in individual 3cm dishes and 500µL of cell suspension (240 000 cells/ml) was spread on top of the slide. The cells were left to adhere for 3h in the incubator before adjusting the amount of medium to 1.5 ml.

### Immunofluorescence

In order to ensure a homogenous staining throughout the thickest colonies, we adapted the procedure described in (Weiswald 2010). Briefly, samples were fixed in a mixture of 4%PFA and 1% Triton in PBS for 1h at 4°C. Then PFA was quenched with a solution of 50mM of NH4Cl and the sample was incubated for 4h in 3%BSA in PBS at 4°C. Antibodies were incubated overnight at 4°C. Nuclei were counterstained with Dapi (molecular probes) and actin with phalloidin 633 (Fluoprobes). Finally, samples were mounted in ProlongGold (Invitrogen). Antibodies: anti-Brachyury (R&D AF2085); Anti-Oct4 (Santa-Cruz sc-9081); Anti-Ki67 (Dako M7248).

## Supplementary Figures

**Movie S1: Time laspe showing ESC colonising a micropattern (Ellipse)**

This movie shows how multiple aggregates of ESC initially form on a micropattern patch before fusing to fully cover the entire shape. Time frames were acquired every 3 min from 4h after plating over a period of 24h.

**Movie S2: Time lapse showing the movement of an ESC colony movements as a cohesive unit**

This movie represents an ESC colony imaged every 3min over a period of 10h. Green cells were cultured at low density in order to enrich this population with T+ cells (∼ 25%). These cells were then mixed to other ESC cultured at control density in order to be able to follow differences in behaviour. This movie shows how, green cells guide the movement of the colony. Please note the extrusion of a green cell trapped in the middle of the colony between 7h and 7h30.

**Fig. S1:**
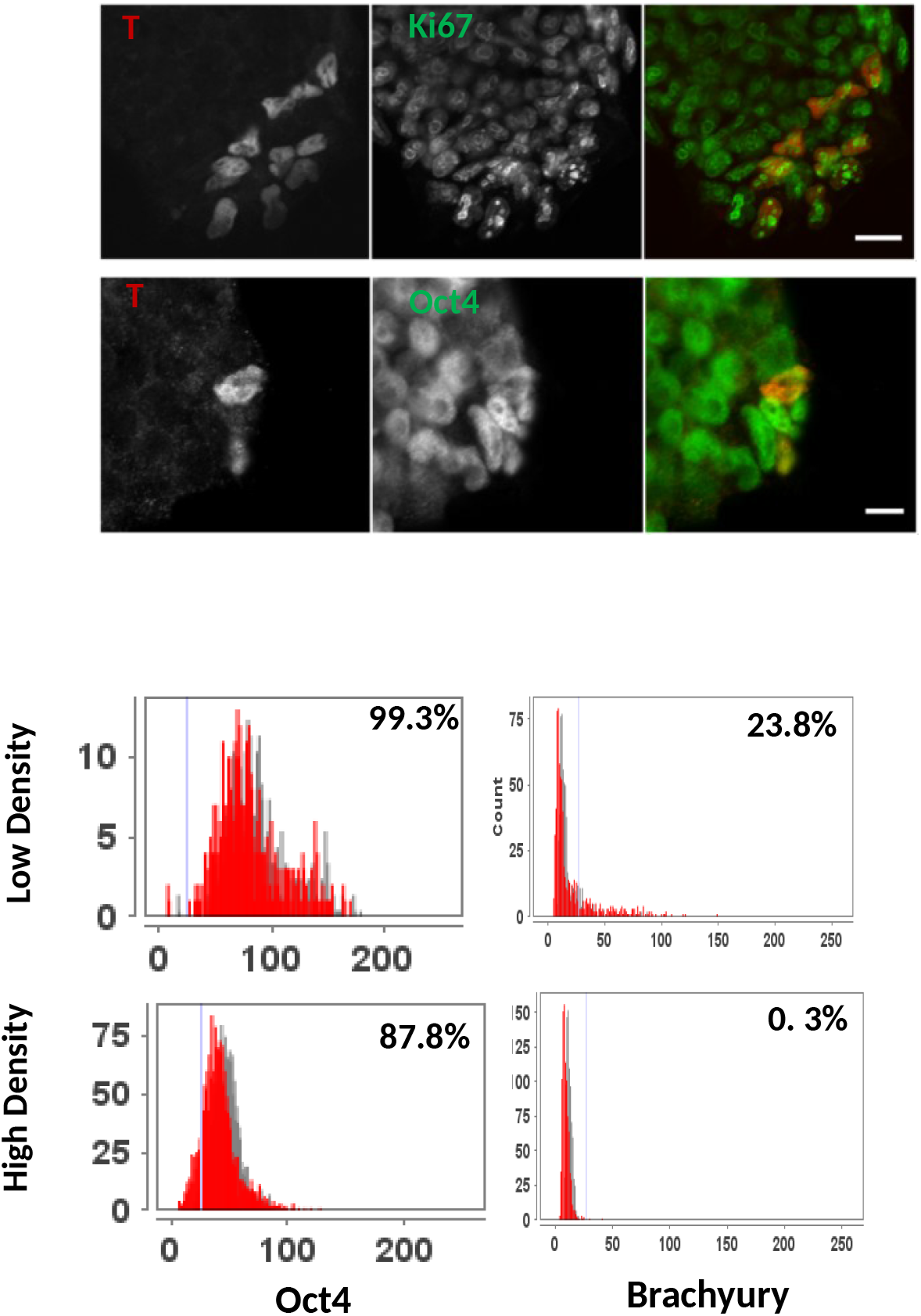
T+ cells detected in Lif/FCS culture are Oct4+ and proliferative consistently with a PS identity. Upper images show the colocalization of nuclear T with the proliferation marker KI67 antigen, (scale bar: 20µm) or with the pluripotency marker Oct4 (scale bar: 10µm). Histograms represent a quantification of the number of T+ cells in ESC populations grown for 48h at low density or at high density.

**Fig. S2:**
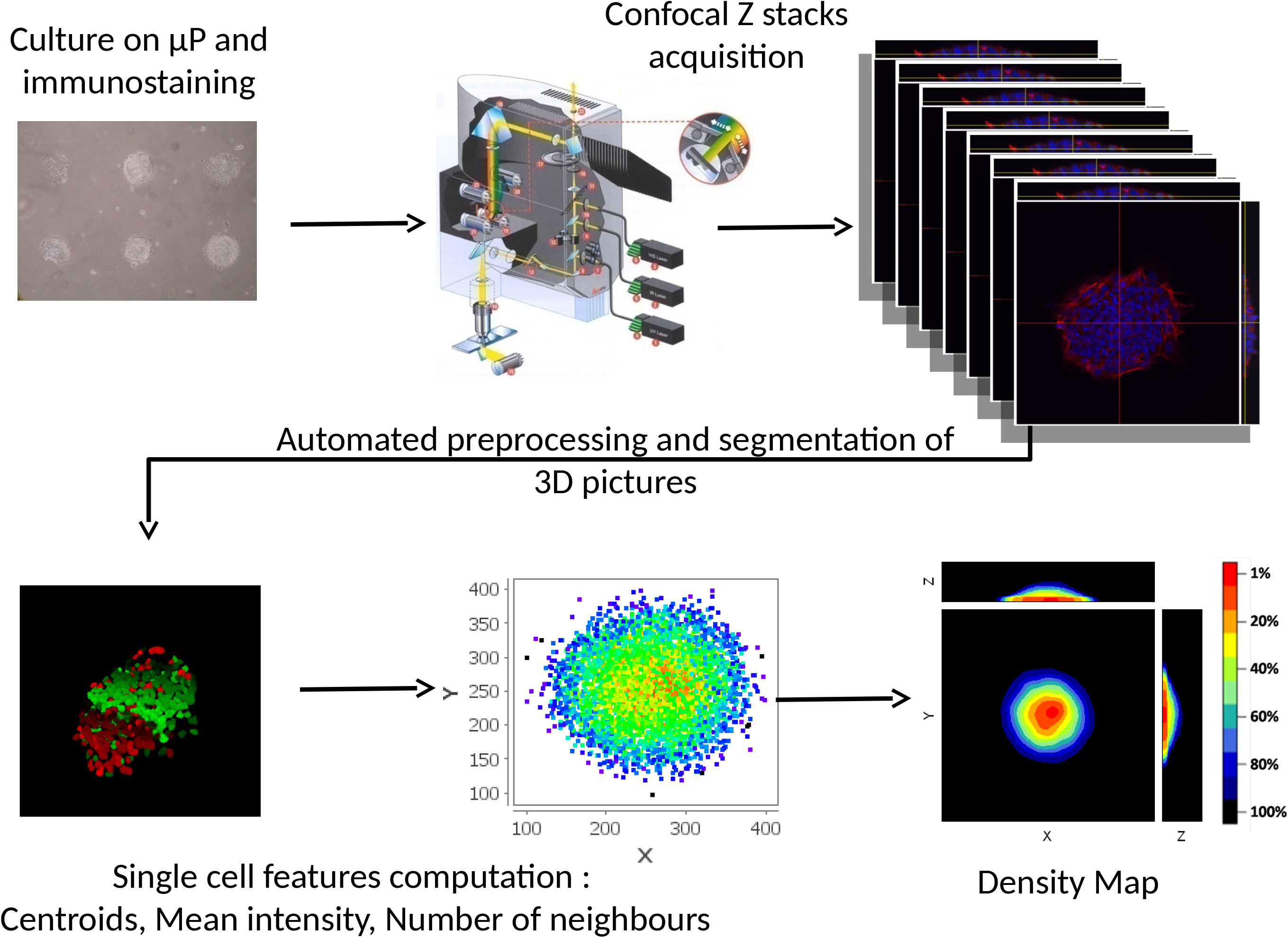
Image acquisition and method for PDM determination. Fixed samples were inspected manually and colonies which fully covered the micropattern and containing at least one T+ cell were selected for imaging. 8-bits 3D images were acquired through an 40X APO objective (NA 1.25) on a Leica SP2 confocal microscope (voxel size: 732 × 732 × 700nm). Channels were acquired sequentially starting with the highest wavelength. We verified that no photobleaching occurred during acquisition by monitoring the level of fluorescence in the z axis. When all 3D images were acquired, each z-stack was registered for the subsequent automated preprocessing, segmentation and analysis (Fig. S1). The preprocessing step consisted in a background subtraction, a gamma correction of the Dapi channel to reveal low intensity nuclei, and a rotation when this was required. To detect cell nuclei in 3D, the dapi signal was segmented using a gradient flow tracking algorithm (Li et al., 2007). Parameters values: sigma = 0.65, minimum nucleus size = 500 pixels and fusion threshold = 1. Analyses were performed with an in-house built java application. The position of each cell was recorded as the centroid of the corresponding segmented nucleus. Each centroid was then tagged with the cell phenotype which was assessed by measuring the average signal intensity in the superimposed area in the other channels of the picture. Cells were considered positive for the nuclear marker of interest when the average pixel intensity was over a threshold determined with a negative control (colony with no T+ cell or Retinoic acid treated cells for Oct4). To generate the probabilistic density maps (PDM), we took advantage of the fact that µP enable the standardisation of the size and geometry of ESC colonies. The colonies were registered in the 3 dimensions by aligning their barycenter computed from the cells centroid coordinates. Finally, the list of cell coordinates with the tag of interest was generated and plotted in R (www.r-project.org/). For PDM representation we used the ks library (Tarn Duong, 2007)to generate the kernel density estimate (KDE) of the distribution. Each PDM presented in the present work is a combination of at least 3 independent experiments. Similarly, for the nearest neighbors analysis, cells were tagged with the number of neighboring centroids within a 30µm radius sphere and the list was plotted as box plots using sigmaplot. For time lapse acquisition, micropatterned slides were fixed under a 3cm dish (MatTek) with grease and tape. Then a mixture of CFSE stained and unstained cells were seeded in the dish 24h prior imaging every 3 min with a Nikon Biostation (x 20 objective). ImageJ plugins: Bio-Formats, Manual TrackJ and ImageJ 3D Viewer which can be found at http://rsbweb.nih.gov/ij/ as well as java libraries: Jfreechart (http://www.jfree.org/jfreechart/) and ujmp (http://www.ujmp.org/) have been used in this work.

**Fig. S3:**
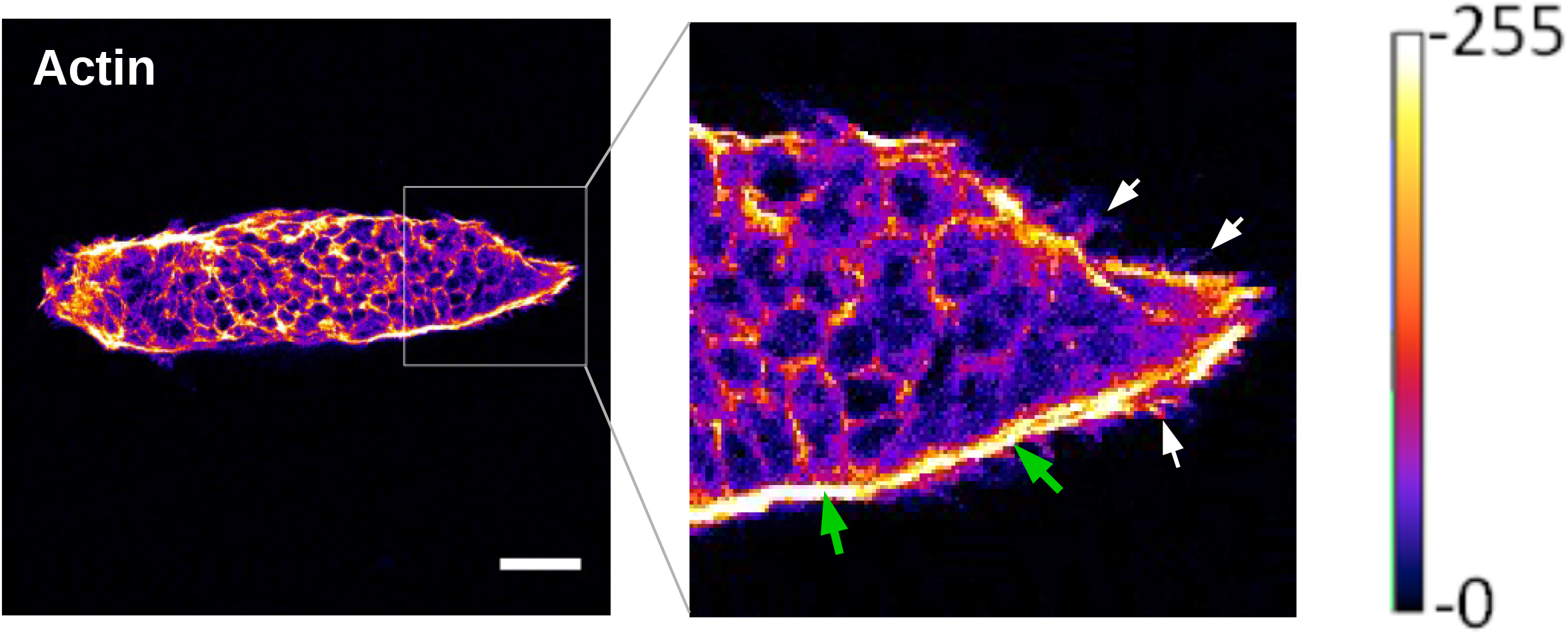
Formation of multicellular actin cables on ellispoidal micropattern. Representative confocal image of an ESC colony grown on an ellipsoidal µP and stained with phalloidin-633. A lookup table has been applied to the 8-bit image (color scale on the right) in order to highlight the actin organization at the multicellular scale. The green arrow points to the presence of a supracellular actin fiber lining the border of the micropattern. White arrows show cell protrusions aligning with the putative distribution of forces within the group. Scale Bar: 50µm.

**Fig. S4:**
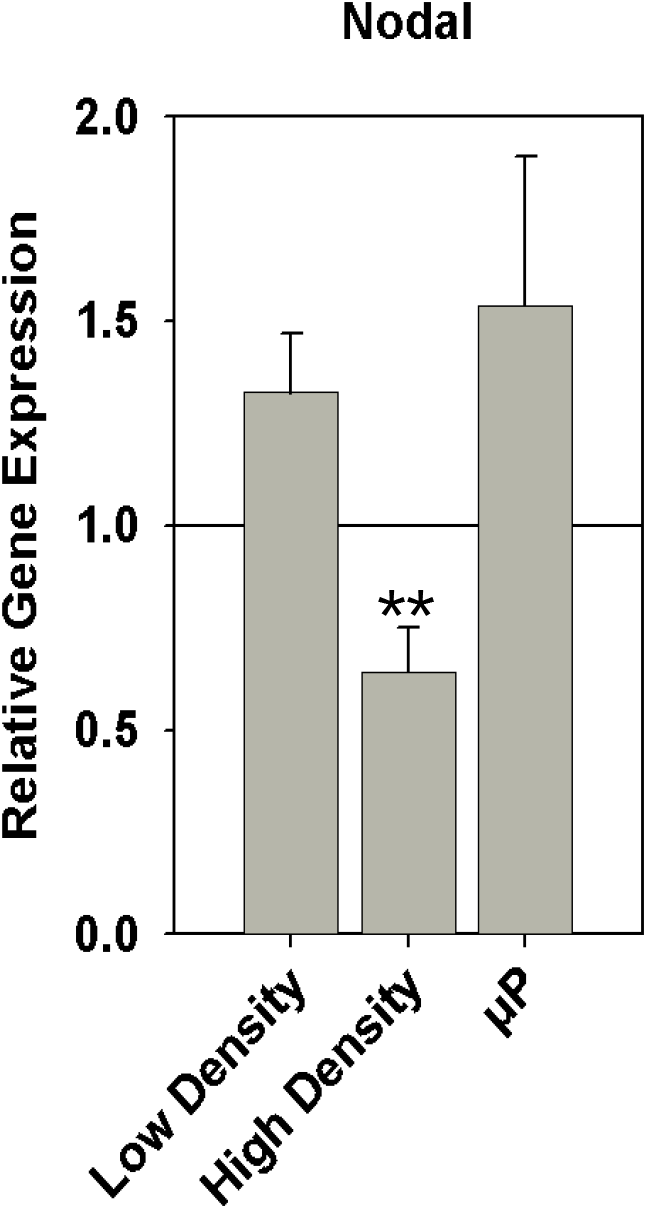
Nodal transcripts levels are lower at high density. qPCR analysis of Nodal expression in cells grown at different densities or on µP. Error bars represent the S.E.M of at least 3 independent experiments. The Student t-test indicate that the difference between high density and low density is significative (** p < 0.01).

**Fig. S5:**
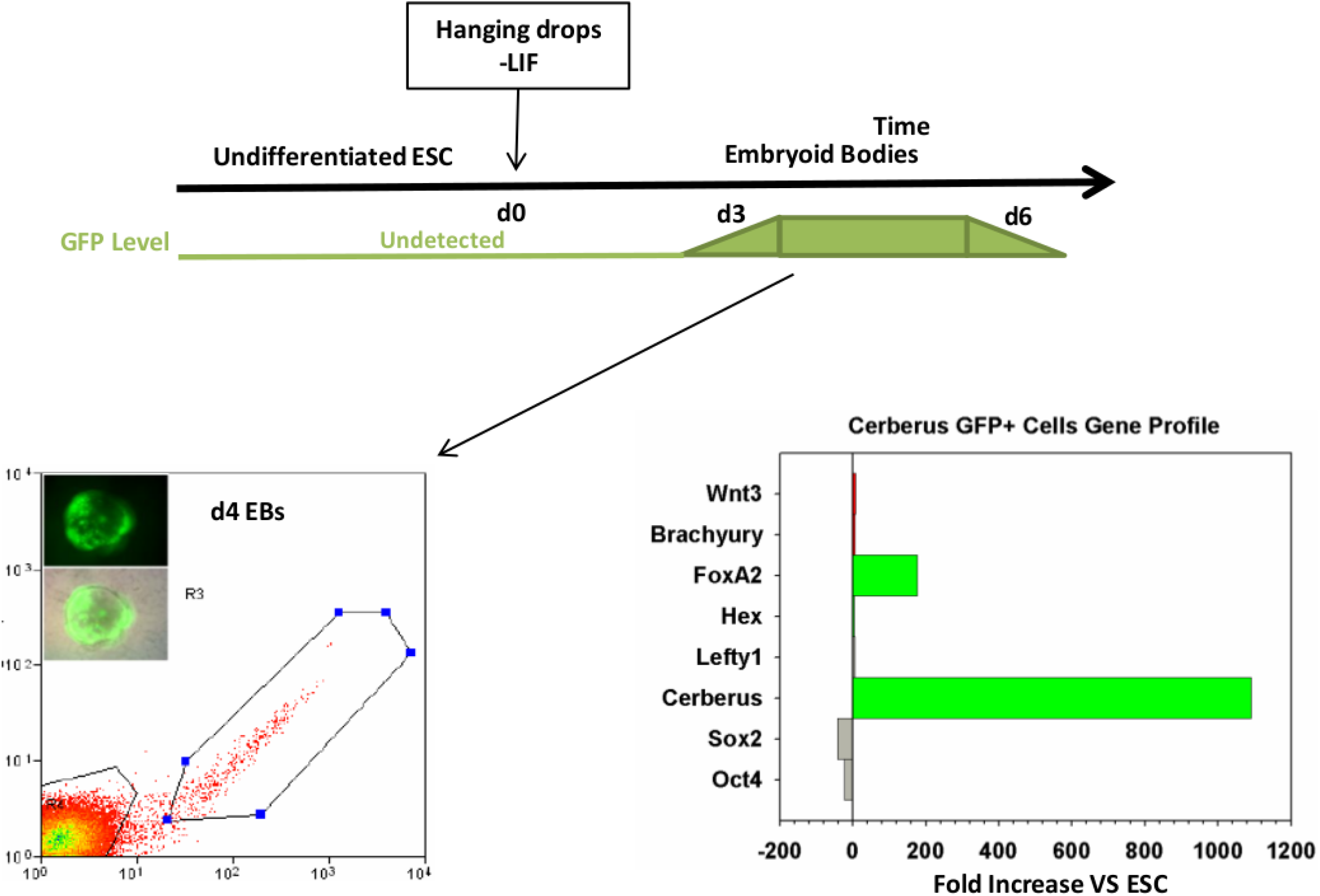
Evidence of Cerberus GFP+ cells in mESC EBs. The upper schematic represents the kinetics of Cerberus GFP expression during EB formation. The bottom panel shows a scatter plot representing the FACS profile of the day 4 sorted GFP+ cell populations. On the right, the bar chart depicts the relative gene expression profile of GFP+ cells vs undifferentiated ESC. NB: This experiment has only been performed once.

